# Replication-associated inversions are the dominant form of bacterial chromosome structural variation

**DOI:** 10.1101/2022.02.15.480581

**Authors:** Matthew D’Iorio, Ken Dewar

## Abstract

Structural arrangement of a bacterial chromosome varies widely between closely related species and can result in significant phenotypic outcomes. The appearance of large-scale chromosomal inversions that are symmetric relative to the *dnaA* gene (usually linked to *oriC*, the origin of replication) has been previously observed; however, the overall prevalence of replication-associated structural rearrangements (RASRs) in bacteria and their causal mechanisms are currently unknown. The decreased cost of full-length genome sequencing has led to a rapidly growing collection of complete genomes spanning multiple different clades, therefore allowing an opportunity to examine chromosomal inversions in the context of species spanning diverse phylogenetic classifications. Here we systematically identify the locations of large, chromosomal inversions in species with multiple complete sequenced genomes using the Refseq and Genbank NCBI databases to investigate potential mediating biological mechanisms. Out of the 239 species available with 10 or more complete genomes, 206 contained sequences with at least one large (≥50Kb) inversion in their set of within-species sequence comparisons. We observed 73.4% of the 127,161 large inversions were centered at a point within 10% proportionate distance to the annotated *dnaA* gene, which is often nearby the origin of replication. Inversions offset from the annotated *dnaA* sequence were generally confirmed to be centered on the actual origin of replication. Equidistant breakpoints from the replication origin and prevalence of flanking repeats provide evidence that the breaks that are formed during the replication process are then repaired to opposing positions. We also found a strong relationship between the later stages of replication and the range in variation of distance from symmetry, suggesting that replication fork arrest may be a mechanistic cause for the asymmetry in some inversions.

## Introduction

Bacterial chromosomes are continuously reorganized by an interplay of mutations, lateral transfer, and mobile genetic elements. Mutations are often broadly categorized as either local – a point mutation affecting a single nucleotide, or global – a structural variant (SV) or rearrangement of an entire locus in the form of an inversion, duplication, insertion, deletion, or translocation. Local mutations are more readily identifiable since they can be detected by comparing relatively short DNA sequence segments, while the discovery of larger structural variants often requires longer draft genomes or complete genome sequences to identify and categorize the full extent of the polymorphism (Periwal and Scaria 2014).

Completed genome assemblies have become fundamental research tools throughout the life sciences. Spurred by multiple generations of massively parallel DNA sequencing technologies, complete genome sequences provide the framework for mutation discovery and can elucidate the mechanisms that influence bacterial evolution. Over the last decade, the continually increasing access to complete genomes offers an unparalleled opportunity to study structural variation across phylogenies and further elucidate its significance in modulating genotype and phenotype variation. Identifying and documenting SVs has been recognized as a promising method for tracking bacterial evolution and providing evidence for the underlying mechanisms that drive recombination (Noureen et al. 2019; Weigand et al. 2019).

Chromosomal rearrangements often result from homologous recombination, which is the process that contributes to the repair of DNA damage and resolves breakdowns that occur during replication (Kuzminov 1999; Michel et al. 2007). Circular bacterial chromosomes generally replicate using replication forks that move bidirectionally away from the replication origin and conclude by meeting at the terminus. Either or both replication forks can break down during regular growth conditions, and their resolution is essential for survival (Cox et al. 2000). A breakdown in both replisomes that is resolved by recombination at a single complex would result in an inversion (Makino and Suzuki 2001), and if both replisomes are moving at the same speed, the inversion will appear symmetric relative to the origin of replication (**Figure 1**). These symmetric, “X”-shaped inversions are henceforth referred to as replication-associated structural rearrangements (RASRs).

**Figure 1.**
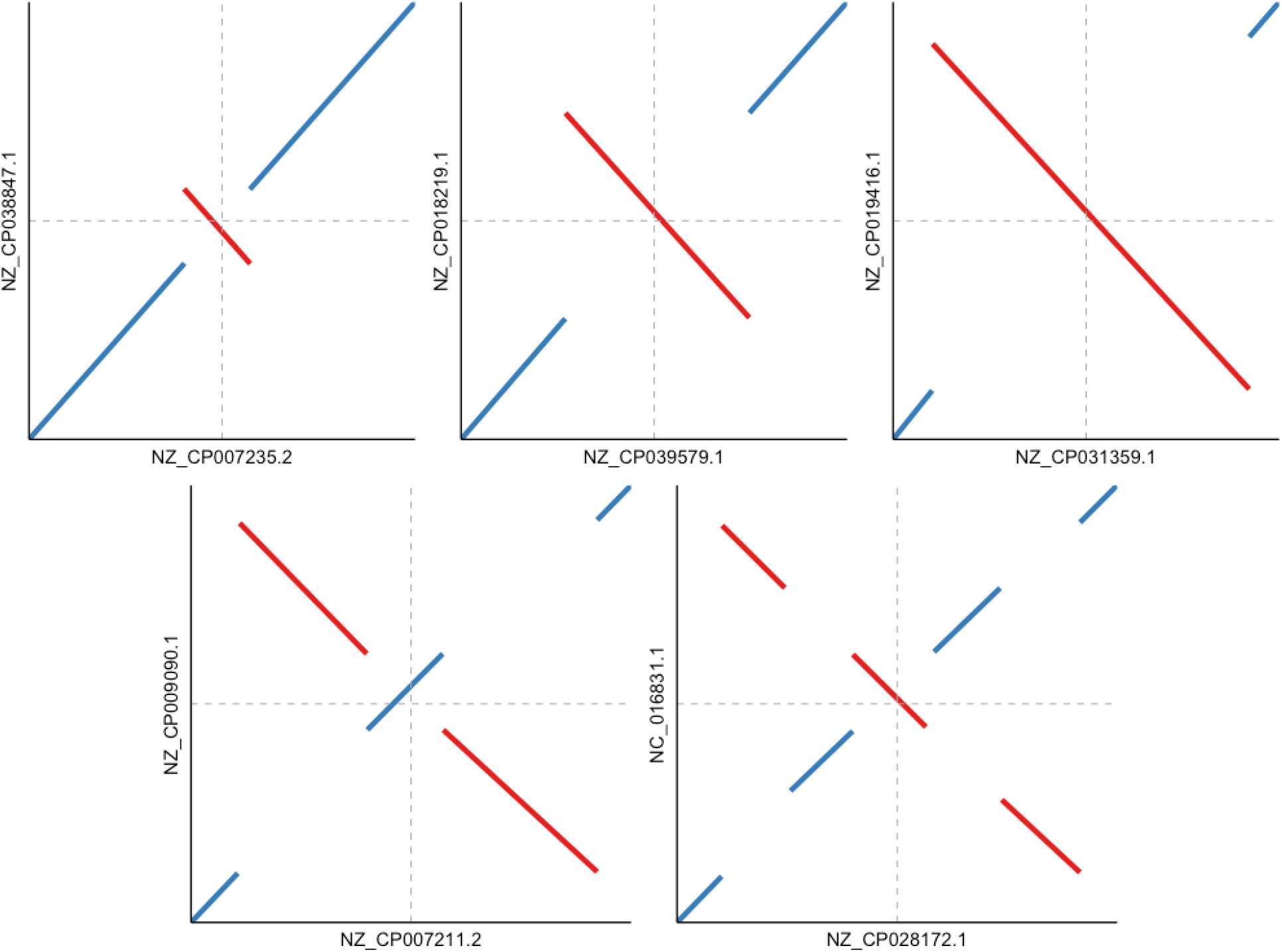
Pairwise comparisons of *Salmonella enterica* sequences showing examples of replication-associated structural rearrangements (RASRs). Each reference is plotted on the X-axis and the query is plotted on the Y-axis. The horizontal and vertical arrows indicate the approximate position of the *dnaA* gene for the query and reference sequence respectively. The top panel illustrates single RASRs of different lengths, and the bottom panel shows examples of double and triple RASRs.

RASRs are highly prevalent forms of SV in well-studied bacteria and were first identified by studying the conservation of the order of genes and the effects of disruptions to conserved gene order (Sanderson and Hall 1970). The significance of RASRs has been established due to their ability to alter phenotypic outcomes such as virulence, survivability, and pathogenicity of eukaryotic hosts (Hughes 2000; Merrikh and Merrikh 2018). For instance, a RASR was observed in isolated colonies of *Staphylococcus aureus*, which regulated the expression of multiple genes and caused phenotypic changes such as antibiotic and immune resistance, growth rate, and colony morphology (Cui et al. 2012). Another RASR spanning 1 Mb in a pathogenic line of *Streptococcus pyogenes* was identified at 40% higher frequency in patient screenings after 5 years, and correlated with a resurgence in severe infections in Japan (Nakagawa 2003). Instances of RASRs have also been extensively documented in model pathogens such as *Salmonella enterica, Escherichia coli*, and *Yersinia pestis* (Roth et al. 1996; Darling et al. 2008). Although synteny is disrupted by these mutational events, the distance of each gene to the origin of replication remains well conserved (Eisen et al. 2000; Tillier and Collins 2000; Repar and Warnecke 2017).

In this study, we have used the NCBI Microbial Genomes Database (Clark et al. 2016 (O’Leary, 2016 #66)) to assess the amount of large structural variation present throughout bacterial phylogeny. Although we restricted our analyses to complete genomes (21,856), which represent only ∼3% of the total number of sequenced bacterial genomes (821,958), it still allowed a survey of 239 species collectively representing the major bacterial clades. Notwithstanding that the databases are heavily skewed towards human and animal pathogens; we have still been able to survey across a wide phylogenetic distribution and a wide variety of ecological niches. Our results confirm previous species-level observations that symmetric inversions represent the predominant form of structural variation. The investigation of breakpoint sequences shows that these inversions are mediated by homologous recombination due to repetitive elements, and the stage of replication influences the degree of symmetry. These RASRs can occur iteratively in successive bacterial lineages, thus offering a potential additional epidemiological tool for tracking strain evolution and distribution.

## Results

### RASRs are identified across phylogenies

We assessed a total of 239 species that were represented by 10 or more complete genome sequences in the NCBI database and identified distinct structural rearrangements in pairwise comparisons of samples within species groups. When two genomes of the same species shared high sequence similarity and did not contain any large structural rearrangements, they were clustered into colinear groups, which we refer to as genoforms. A genoform in this work refers to a group of collinear chromosomal sequences within a species with at least 80% sequence similarity and without an inversion spanning more than 50 Kb. This size of inversion represents approximately 1% of the genome length and is clearly detectable in a pairwise comparison segment plot, since genome lengths range within our dataset from 0.4 Mb to 9.8 Mb with an average of 4.7 Mb.

We were able to identify multiple distinct genoforms in 84% of species included in our dataset across 9 major clades. The number of RASRs and distinct collinear clusters identified generally increased with the amount of sequence data available for each species, and some species appeared to have a higher propensity for incurring structural variation than others. Some species groups contained many fully sequenced genomes and little evidence of structural variation such as *Mycoplasma pneumonae* (78 genomes, 1 genoform) and *Chlamydia trachomatis* (86 genomes, 1 genoform). Genome stability for both species could be related to their slower reproduction rates relative to other bacteria (Vieira-Silva and Rocha 2010). In contrast, some species appeared to have a higher propensity for rearrangements such as *Yersinia pestis* (58 genomes, 46 genoforms), *Pseudomonas putida* (42 genomes, 29 genoforms) and *Pseudomonas syringae* (36 genomes, 23 genoforms); all of which have relatively fast reproduction rates and higher repetitive genome content compared to other species.

Although our data was dominated by mammalian pathogens, we were also able to identify RASRs in bacteria that occupy a range of biological niches. There were instances of RASRs in different probiotic species such as in 3 out of 4 *Bifidobacterium* species, 7 out of 9 *Lactobacillus* species, and *Streptococcus thermophilis* and S. *salivarius*. We also found symmetric inversions in bacteria from environmental samples and plant symbiots such as *Bradyrhizobium diazoefficiens, Phaeobacter inhibens*, and *Rhodopseudomonas palustris*.

Incidences of RASRs were observed throughout the major clades of our dataset. We adjusted for the over-representation of Pseudomonadota (previously Proteobacteria, (Oren and Garrity 2021)) by partitioning their species at the class level to characterize occurrence across the major clades. We observed that these types of inversions can be iterative, leading to increasingly complex genome shuffling in successive strain lineages (**Figure 1**), and examples of RASRs across bacterial phylogeny are displayed in **Figure 2**. In this report, we now use nomenclature which The International Committee on Systematics of Prokaryotes (ICSP) recently implemented to revise prokaryotic phylum names (Whitman et al. 2018; Oren and Garrity 2021), but also display the previous nomenclature in brackets for each clade that has changed.

**Figure 2.**
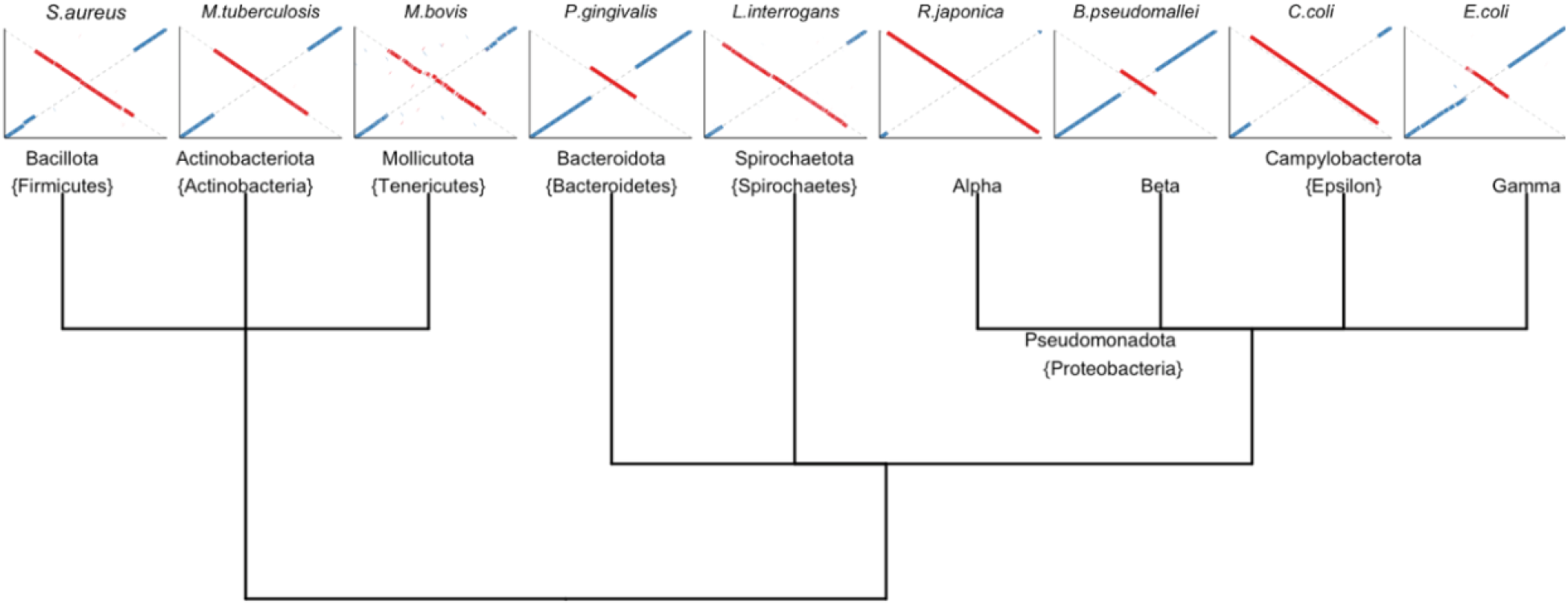
Nine phylogenetic clades containing species with at least one large-scale structural variant. RASRs were extensively documented throughout clades, and an example of a representative organism from each is displayed below each label.

### RASRs are the predominant form of chromosome structural rearrangement

The symmetry of each inversion was evaluated as the proportionate distance from the *dnaA* gene set to position zero (analogous to 12 o’clock), where a value of zero would indicate maximum symmetry, and positive and negative values indicate proportionate distance away from *dnaA* to the left and right respectively. The locus for the terminus is directly opposite of the origin for most bacterial species (analogous to 6 o’clock). Out of an observed 127,161 inversions between all genoform comparisons within our species groups, 57.7% of inversions were less than a 0.05 proportional distance from symmetry, and 73.4% were within a 0.1 proportional distance from symmetry. The tendency towards symmetry in the full dataset was also indicated by the overall average the proportionate distance being 0.001 and a 95% CI [-0.011, 0.013]. Our dataset containing all inversions was overrepresented by species that contained disproportionately high counts of inversions such as E. *coli* (47,973 inversions), *Bordetella Pertussis* (14,653 inversions), and Y. *pestis* (10,416 inversions). These were the top three most represented genomes and together comprised 57% of the total count of inversions throughout the dataset. To identify the tendency of inversions throughout our 193 species, we recorded the average inversion midpoint value for each species and plotted the distribution on a circular representation of the bacterial chromosome (**Figure 3**). The average midpoint of an inversion was calculated as the mean between the central point of each inversion on the coordinates for both the reference and the query genomes. We used proportionate midpoint values as a relative value for genome length to normalize for the wide range of genome sizes.

**Figure 3.**
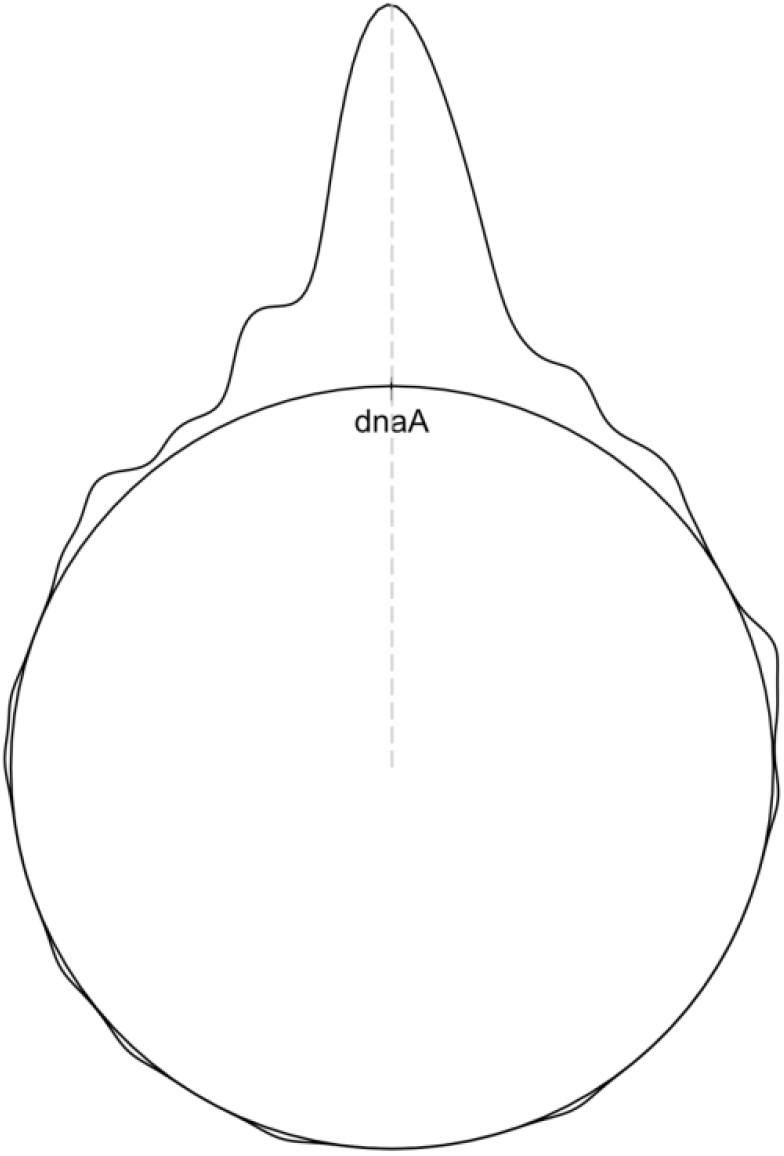
The aggregate distribution of average midpoint values for all large-scale inversions in each of the 196 bacterial species.

### Variation in RASRs across species groups

To understand the overall distribution of inversion symmetry, it was informative to normalize for genome length and calculate the distance as a metric to the left and right of the origin as well as the absolute distance from symmetry. The distribution of symmetry distances within species indicates which species conform to or deviate from the overall trend of symmetry. Using the counts of inversions in species with at least 20 inversions, we identified the distribution of symmetry distance in 101 species (**Figure 4**). Although the mode of distance for most species was at or very close to zero, we identified multiple species with bimodal distributions that were significantly offset from symmetry. In Gammaproteobacteria, the species *Haemophilus parainfluenzae, Manheimia haemolytica*, and *Haemophilus ducrei* all contained inversions that were predominantly located away from the replication origin (**Figure 4a**). We also observed species with a single node that was consistently offset from symmetry in one direction such as in *Bordetella pertussis* and *Riemerella anatipestifer* (**Figure 4b**). Since the *dnaA* gene is not always adjacent to the origin of replication, we identified the minimum distance between *dnaA* and the origin of replication annotated using the Z-curve method from the Doric database (Luo and Gao 2019). Three possible *dnaA* to OriC distances were recorded for each species: (1) average negative values to represent OriC to the left of *dnaA*, (2) average positive values to represent OriC to the right of *dnaA*, and (3) values of zero to represent *dnaA* as directly beside *dnaA*. The distance from *dnaA* to OriC roughly corresponds to the distance we observed between *dnaA* and the mode of inversion midpoints.

**Figure 4.**
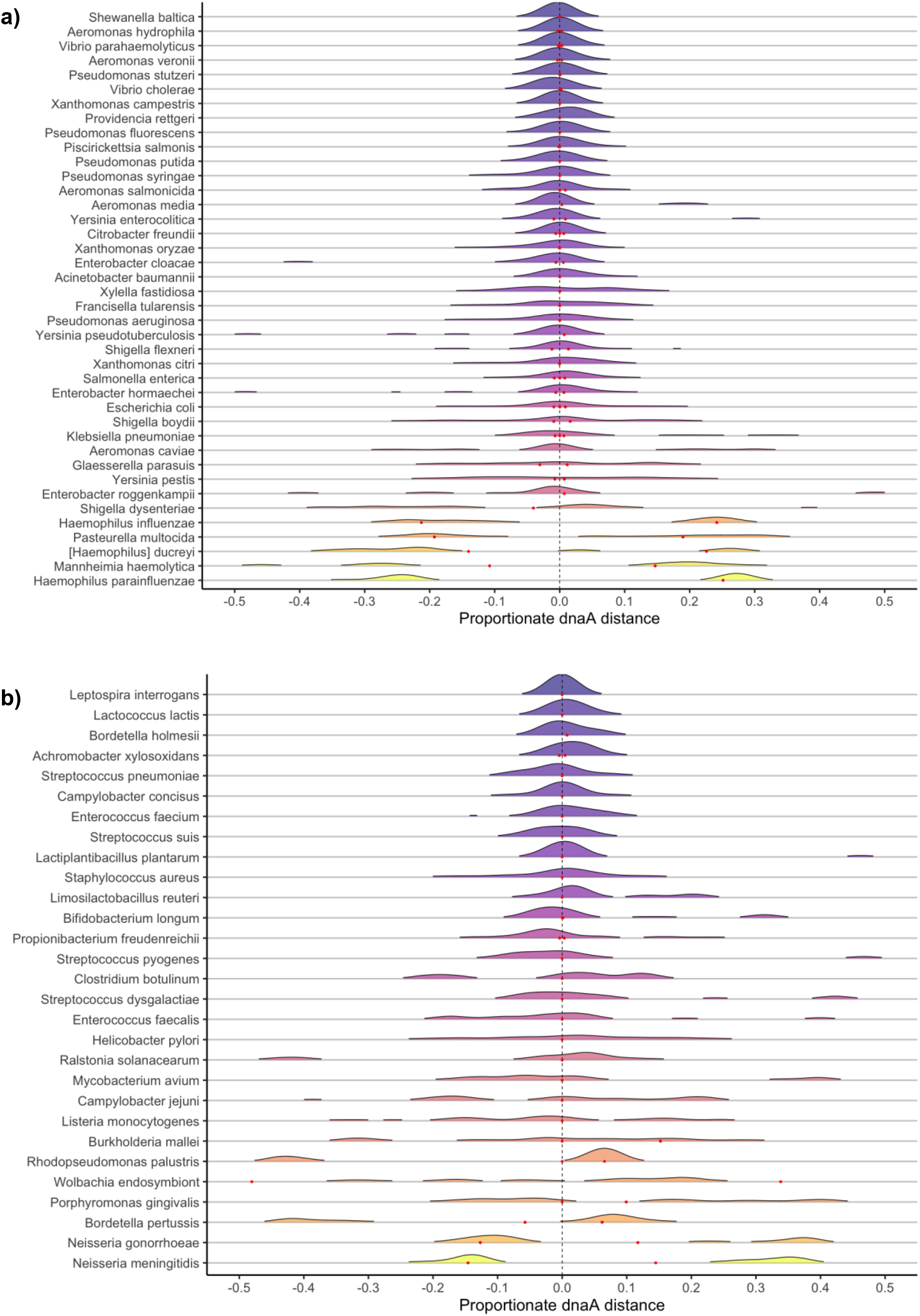
The distribution of midpoints for all Gammaproteobacterial species (a) and all other species (b). Point 0 represents the position of the annotated *dnaA* gene. The red points indicate the approximate positions of the origin predicted by the doric database.

### RASRs are mediated by homologous recombination during replication

Replication and recombination are interdependent processes of DNA metabolism (Syeda et al. 2014). While replication forks move bidirectionally and independently along each side of the chromosome, the progression can be halted and the RecBCD pathway can initiate fork reversal to restart replication (Michel et al. 2004). Interspersed repeated segments may be mediators in the process of forming RASRs given that an arrest in replication could become resolved by reinitiating at opposite forks that share the same sequence. Repeated elements within the chromosome have been found to mediate chromosomal structural recombination (Achaz et al. 2003).

To identify the association between repeat content and inversion occurrence, we calculated the mean proportion of repeat content for each species and compared that to a standardized count of inversions per comparison We found a low but significant overall association between repeat content and inversions per comparison using linear regression (**Figure 5a)**. The role of repeat-induced mediation of inversions was further examined by extracting the overlapping sequences spanning the breakpoints of the inversions. As an example, the pairwise comparison of two sequences within *Erwinia amylovara* produced a single inversion with overlapping segments spanning each breakpoint (**Figure 5b**). The two breakpoints for the inversion overlap with roughly 5.6kb of colinear sequence on both sides, and both breakpoint sequences share nearly identical nucleotide similarity and the same collection of genes. The 5.6 Kb sequence at each breakpoint contains 4 genes and the complete sequence was identified only at the breakpoint loci for both genomes (**Figure 5c)**. Partial repeats of the breakpoint sequence were also found in two other loci in both genomes, but no rearrangements using these loci have been observed.

**Figure 5.**
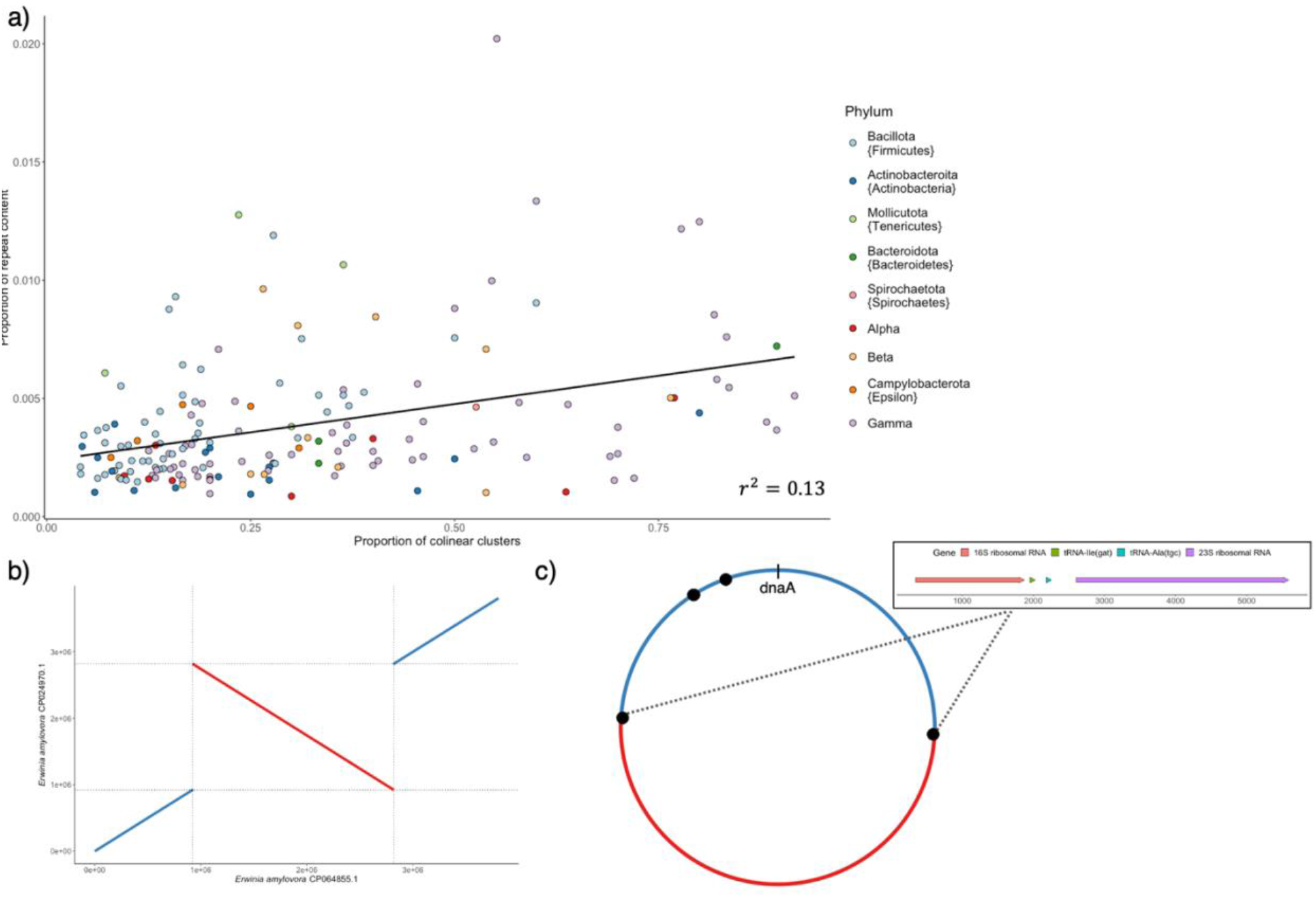
Relationships between repeat content and large structural rearrangements; (a) The mean proportion of repeated 31-nucleotide bocks (k-mers) in a whole-genome plotted against the number of genoforms (distinct colinear clusters) within each species group, (b) two genomes of *Erwinia amylovara* that with a single inversion containing identical overlapping breakpoints at the ends of the inversion, and (c) A circularized representation aligning the two *Erwinia amylovara* genomes highlighting the annotated repeat sequence found at each breakpoint; this sequence is only partially repeated at only two other loci within both genomes (marked by the two points appearing to the left of dnaA).

### De-synchronization causing asymmetry in inversion formation

The independence of replication forks is exemplified by the observations that unequal replisome distance from the origin is commonly observed throughout the replication process and that one replication fork may be halted while the other continues (Breier et al. 2005). Fluorescent labelling of live *E. coli* cells has provided evidence that sister replichores operate independently and the distance between them varies throughout the process of replication (Reyes-Lamothe et al. 2008). If large-scale chromosomal inversions occur within progenitor strands during replication, then an inversion that is offset from symmetry could be caused by an unequal speed of synthesis between replication forks. The offset distance relative to the *dnaA* locus caused by unequal replication time would be exacerbated towards the end of replication. Using the length of individual inversions as an index for the approximate replication time, we evaluated the relationship between replication length and the proportionate distance from *dnaA* (**Figure 6**). The overall trend was heteroskedastic with roughly equal counts of inversions on either side of *dnaA*; 49% were negative (offset to the left) and 51% were positive (offset to the right). To compare the variation in values over replication time, the values were sorted by time and split into 42 groups with equal counts of distance measurements and the maximum and minimum values were calculated from each group. The linear model was then generated comparing the maximum and minimum value groups with the average replication index for each group of observations, where maximum values correspond to distance from *dnaA* to the right, and minimum correspond to distance to the left. The average time against maximum distance corresponded with further distance over the proportionate replication index at a rate of 1.53 (r^2^=0.63, p<<0.01), while the average time against the minimum distance decreased at a rate of 1.34 (r^2^=0.64, p<<0.01).

**Figure 6.**
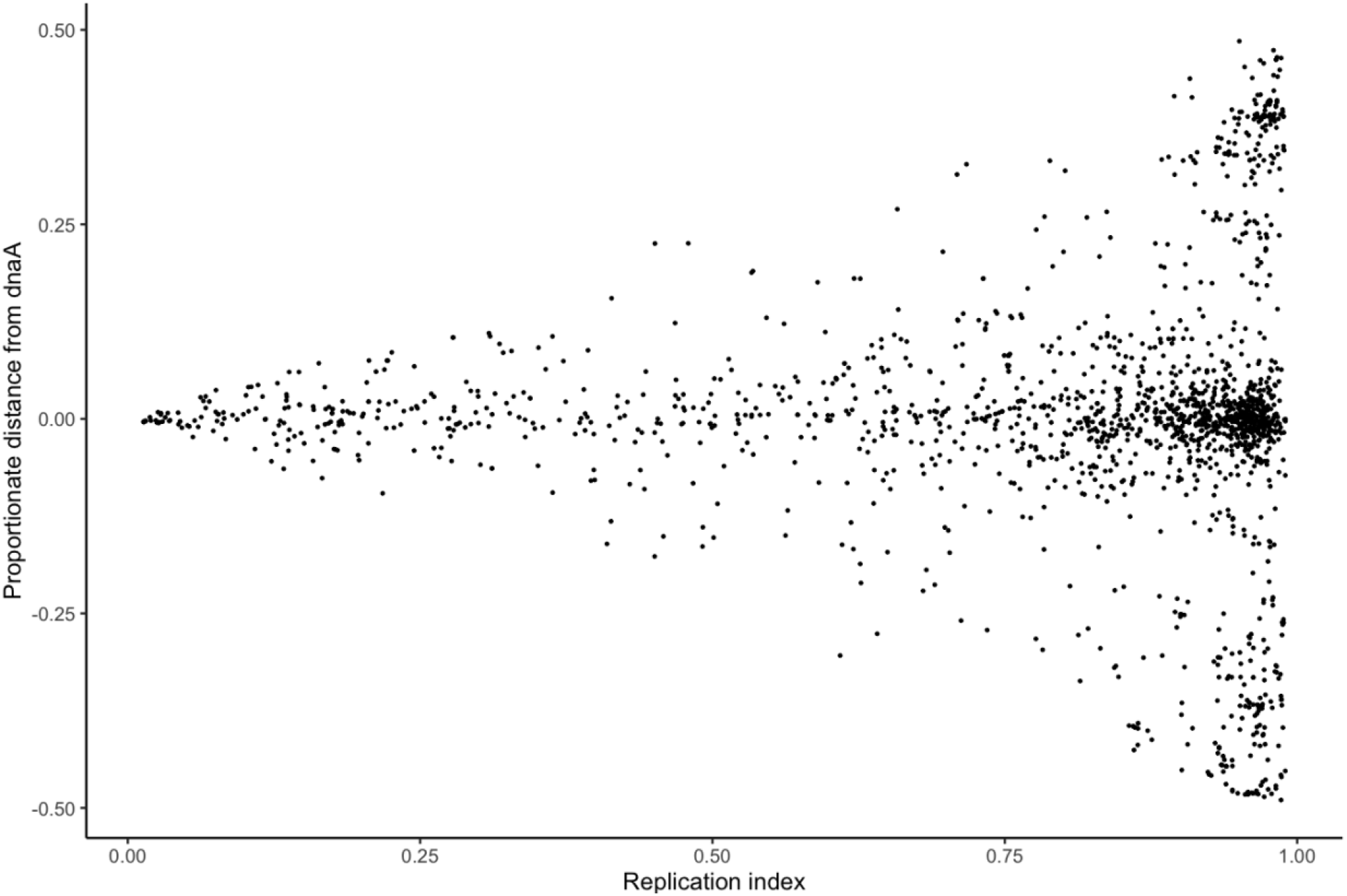
The proportionate distance from *dnaA* compared to the replication index for all individual inversions. The replication index reflects the span of newly synthesized chromosomes when the inversion occurred.

## Discussion

In this paper, we have investigated the prevalence and potential mechanisms of chromosomal inversions symmetric to the origin of replication using NCBI’s full collection of complete genome sequences. The overall prevalence of RASRs is evident from the high frequency of inversions that are observed to be centered near or directly on the origin of replication. The predisposition for structural variation can be influenced by the proportion of repeat content, given the evidence that an interspersed repeat was often found at inversion breakpoints. A strong tendency for inversions to congregate around the origin of replication is consistent with previous models that have suggested that inversions can be the result of breaks that form during the replication process. While the overall prevalence of repeat content was not always associated with a higher incidence of inverted segments, equally interspaced repeated sequences appear to be an important component of the complete mechanism driving structural variation. Given the overall prevalence and the iterative patterns observed between comparisons, this feature could be integrated into phylogenetic analyses as an evolutionary distance metric.

We also note multiple lines of evidence excluding the possibility that the appearance of large-scale symmetrical inversions is simply a result of technical errors in DNA sequencing or computational assembly. First, given the wide range of species, sequencing centers, sequencing dates, and sequencing strategies that have been used, this consistent pattern of genetic diversity is highly unlikely to be the result of technological errors. Second, the observation of having these inversions centered on *dnaA* indicates coordination of breakpoints, again making a sequencing error unlikely. Third, in instances where we can recover genome assemblies and the long reads underlying them, we can conclusively show that multiple independent reads clearly demonstrate that the inversion is a biological difference and not an assembly artifact. Conservation of inversion sizes and evidence of iterations over generations suggests that the propensity for symmetric chromosomal inversions is a basic form of genetic variation common to all types of bacteria.

The strong overall tendency for inversions to be symmetric to the origin of replication provides evidence that the underlying mechanisms are driven by breakage and misrepair during the replication process. DNA supercoiling before and after replication forks may be the cause in the formation of a single complex wherein crossing over of replichores along with double-ended breakage and repair ultimately forms the inversion. If both break sites contain identical sequences (i.e. interspersed repeats), then there should be an increased likelihood of misrepair to opposite ends. A single daughter chromosome also has both a leading and lagging strand synthesizing at each replichore, and our observed shifts away from symmetry may be a direct result of the difference in speed of synthesis. Moreover, since the termination site of replication is often but not always 180 degrees away from the origin, the instances where the termination site is asymmetrical could influence the axis of an inversion (Kono et al. 2014). This range of possible errors is exacerbated if the break occurs at a later stage in the replication process. The even range of distance from either side of the replication origin as an overall trend suggests that the error rate in both replication forks move at a similar speed with a comparable error rate.

The pervasiveness and iterative nature of these RASRs underscore the potential for this method of clustering to be used as a method for tracking bacterial evolution. Structural variation analysis could add a useful dimension for phylogenetic analysis that could complement current methods such as nucleotide and gene content similarity. The ability to apply this method across species will be aided by the increasing rate of publishing complete genome sequences. Full-length genomes still represent a very small proportion of publicly available bacterial sequence data given the resources and costs associated, however, their potential to aid in the discovery of the intricacies of bacterial diversity has not yet been fully realized.

## Methods

### Data collection

Sequence data was downloaded directly from the NCBI RefSeq and Genbank public datasets on (26/01/21). All RefSeq and Genbank assembly summaries were combined and filtered to generate a unique table containing all RefSeq and nonredundant Genbank summaries. Species with fewer than 10 available sequences were omitted. The corresponding FASTA and GFF files were downloaded for all other assemblies that included an annotated *dnaA* gene. We also used data from the Doric 10.0 bacterial database (Luo and Gao 2019), which was last updated (17/06/18).

### Synchronization to *dnaA*

NCBI submissions of circular bacterial genomes may not have standardized starting points and orientations. Since the *dnaA* gene is ubiquitous in bacteria and is a marker for the position of the origin of replication, we used it as a starting point for pairwise comparisons of a linearized representation of the genome sequences. We synchronized every genome sequence by using the coordinates of the *dnaA* gene from each assembly GFF file to re-orient and rotate each sequence as necessary to create the same starting position and polarity for every sequence. We developed a standalone webtool to synchronize individual genomes that is freely and publicly available at https://www.computationalgenomics.ca/tools/bacterial-genome-synchronizer/. Genomes that did not have an annotated *dnaA* sequence, and those with multiple *dnaA* sequences, such as those within Chlamydiae, are currently omitted from the pipeline.

### Pairwise alignment and collinearity clustering

After synchronization of each genome, all sequences were pairwise compared to all other members of the set of sequences corresponding to the same species. We used MASH (Ondov et al. 2016) to provide a nucleotide similarity score, and penalized the score when genomes were different lengths by subtracting a normalized value of the sequence length difference. We used this measurement to select the genome that had the highest similarity to all other genomes within the species as the starting point used to cluster for collinearity. This sequence was then used as the reference sequence that was aligned to all other sequences within the species using MUMmer4 (Marçais et al. 2018). Any sequence that did not show an inversion at a minimum length of 50,000 and had at least 80% of the reference covered by alignments was considered collinear. This step is repeated for all non-collinear sequences until each was sorted into a collinear cluster. After sequences were sorted, each cluster was pairwise compared using the two representative sequences that had the highest shared nucleotide identity.

The complete collection of inversions between each combination of clusters was recorded as the reference and query position intervals for each species. Each inversion was summarized by taking the average value of the center of the intervals spanning the inversion for the reference and query sequence and divided by the length of each genome. Distances are measured proportionately from −0.5 to 0.5 representing the distance to the left or right from *dnaA* respectively, and a value at zero represents perfect symmetry. We calculated distributions for the average midpoint for each species, and the totality of inversions within species that contained at least 20 inversions. We also used the average absolute distance for all species to identify the overall distance from symmetry.

We also determined the proportion of each sequence that was repetitive to measure the association between repetitive content on genome stability. This was done by counting the total number of repeated k-mers at a length of 31 nucleotides and dividing by the total number of possible 31-mers based on the full length of the genome. This was used to compare the measurement of repeat content against the prevalence of inversions within a species.

## Acknowledgments

We are indebted to the NCBI microbial genomics databases. We are deeply grateful to the multitudes of sequencing centers and personnel for their efforts in generating high-end genome sequences and their commitment to free and open access data sharing. We thank Professors Greg Marczynski, Mathieu Blanchette, and Rodrigo Reyes-Lamothe for advice, guidance, and critical reviews of earlier versions of this work. MD is supported by the McGill Genome Centre and Genome Canada.

